# Loss of BubR1 Acetylation instigates Replication Stress leading to Complex Chromosomal Rearrangement in tumors

**DOI:** 10.1101/752642

**Authors:** Jiho Park, Song-Yion Yeu, Sangjin Paik, Junyeob Lee, Jinho Jang, Semin Lee, Young-Il Koh, Hyunsook Lee

## Abstract

Chromosome number and structure instability is the hallmark of cancer. Equal chromosome segregation is guaranteed by the spindle assembly checkpoint (SAC), thus defective SAC leads to chromosome instability. However, aneuploidy alone is not oncogenic, and whether compromised SAC is associated with structure instability remains elusive. BubR1 is a core component of SAC, which is acetylated at lysine 250 in mitosis. Previously, we showed that deficiency of BubR1 acetylation in mice (*K243R/+*) leads to spontaneous tumorigenesis via chromosome mis-segregation. Here, we asked whether loss of BubR1 acetylation is associated with chromosome structure instability by examining *K243R/+* mice intercrossed to *p53*-deficient mice. Genome-wide sequencing and spectral karyotyping of the double mutant mouse tumors revealed that BubR1 acetylation deficiency leads to complex chromosome rearrangements, including Robertsonian-like whole-arm translocations and premature sister-chromatid separations (PMSCS). In primary MEFs, replication stress was markedly increased in telomeres and centromeres, suggesting that the replication stress underlies the significant increase of DNA damage and subsequent chromosome rearrangements. Furthermore, defects in BubR1 acetylation at K250 were detected in human cancers as well. Collectively, we propose that chromosome mis-segregation by the loss of BubR1 acetylation causes chromosome structure instability, leading to massive chromosome rearrangements through the induction of replication stress.

## Introduction

Accurate chromosome segregation is guaranteed by the spindle assembly checkpoint (SAC). Therefore, loss or mutation in the checkpoint component was thought to be the cause of aneuploidy and cancer. However, studies in genetically engineered knockout mice revealed that aneuploidy, instability in chromosome numbers, is not oncogenic by itself. Only when combined with genotoxic stress, aneuploidy contributed to tumorigenesis (1).

BubR1 is a core component of the spindle assembly checkpoint (SAC), constituting the mitotic checkpoint complex (MCC) in SAC (2-4). Homozygous deletion of *BubR1* in mice leads to embryonic lethality; heterozygous mice develop megakaryopoiesis with aneuploidy but do not show any signs of spontaneous tumorigenesis (5). Moreover, studies from hypomorphic *BubR1* allele suggested that BubR1 insufficiency is associated with premature aging but not spontaneous tumorigenesis (6).

BubR1 is acetylated at lysine 250 exclusively in mitosis. Acetylation of BubR1 modulates APC/C activity that acetylated BubR1 is the inhibitor of APC/C (7), whereas deacetylation of BubR1 is a cue for SAC silencing and mitotic exit (8). Importantly, K250 locates adjacent to ABBA motif, KEN box, and D-box (4, 9), corroborating that BubR1 acetylation/deacetylation is a molecular switch regulating APC/C (7). Furthermore, the breast cancer susceptibility gene product, BRCA2 mediates BubR1 acetylation as it recruits the acetyltransferase PCAF to BubR1. Therefore, *BRCA2*-deficient cells fail to acetylate BubR1 at K250 (10). Consistently, transgenic mice that were engineered to inhibit BRCA2 and BubR1 association, thus disrupting the acetylation of BubR1, developed tumors. The result suggests that without the perturbance of BRCA2’s role in homology-directed DNA repair, deficiency in regulating BubR1 acetylation can lead to spontaneous tumorigenesis (10).

Congruently, mice heterozygous for BubR1 acetylation at lysine 243 (*K243R/+*), corresponding to K250 in human, develop spontaneous tumors in ∼ 12 months without apparent signs of premature aging (11). In the mouse study, it was found that acetylation of BubR1 at lysine 250 has dual roles: maintenance of MCC in checkpoint signaling and stabilization of chromosome-spindle attachment by recruiting PP2A-B56α subunit for antagonizing Aurora B kinase in chromosome congression (11).

Chromosomes from *K243R/+* MEFs display near-diploid aneuploidy. Also, premature sister chromatid separation (PMSCS), and inter-chromosomal translocation were observed, albeit to a low level. Micronuclei increased significantly, consistent with marked failure in error correction in the chromosome-spindle attachment (11). Furthermore, analysis of *p53* cDNA in primary tumors showed that missense mutations were apparent, just like human cancers, in the tumorigenesis of acetylation-deficient BubR1 mice. These results led us to hypothesize that genetic instability and chromosome structure aberrations may be the underlying mechanism that leads to tumorigenesis in *K243R/+* mice.

Along these lines of thinking, we were prompted to ask if chromosome mis-segregation by the loss of BubR1 acetylation provoked chromosome structure instability, in addition to chromosome number instability, providing the basis of tumorigenesis. For this, *K243R/+* mice were intercrossed to *p53*-null mice to rescue all chromosome aberrations and mutations during proliferation in acetylation-deficient BubR1 mice. As *p53* mutations were frequently found in tumors of *K243R/+* mice (11), the breeding scheme was physiologically justified. We were also interested to see if chromothripsis, one-time chromosome crisis and cis-rearrangement of chromosomes (12,13), was associated with the tumorigenesis of BubR1 acetylation-deficient mice, as micronuclei were suggested source for chromothripsis (14) and *K243R/+* cells exhibit marked increase of micronuclei during proliferation (11).

Here, we show that chromosome mis-segregation by the loss of BubR1 acetylation instigates replication stress and DNA damage, leading to complex chromosome structure instability, including PMSCS, inter- and intra-chromosome rearrangements, and Robertsonian whole-arm translocations. These features were also found in human cancer, suggesting that BubR1 acetylation deficiency is oncogenic, and may be associated with cancers of complex karyotype.

## Results

### Intercross of *K243R/+* mice to *p53*-null mice results in acceleration of tumorigenesis and change in tumor spectrum

Previously, we observed a small fraction of chromosome translocation and missense mutations, in addition to aneuploidy in primary MEFs from acetylation-deficient BubR1 (*K243R/+*) mice (11). As aneuploidy alone is not oncogenic and the outcome of aneuploidy in cell proliferation is complex (1, 15, 16), we reasoned that the chromosome structure aberration and genetic instability in *K243R/+* mice might be the cause of tumorigenesis, while aneuploidy aggravates the situation (11).

To understand the basis of tumorigenesis by mitotic infidelity, we decided to analyze all chromosomal and genetic aberrations in *K243R/+* mice. For this, we crossed *K243R/+* mice to *p53*-null mice to rescue all aberrations in *K243R/+* mice in the road to tumorigenesis. We generated cohorts of WT, *K243R/+, p53-/-*, and *K243R/+;p53-/-* mice and monitored for the development of tumors for up to 80 weeks. *K243R/+* mice develop spontaneous tumors in 12 months with the frequency of ∼ 40% (11). In comparison, all of the examined *K243R/+; p53- /-* mice developed tumors (n=33, 100%), as was *p53-/-* mice (n=14, 100%). The median tumor-free survival time was ∼ 24 weeks in *K243R/+;p53-/-* mice, while it was ∼ 65 weeks in *K243R/+* mice. *p53-/-* mice exhibited similar tumor-free survival time as *K243R/+; p53-/-* (Fig. 1A).

**FIG 1.**
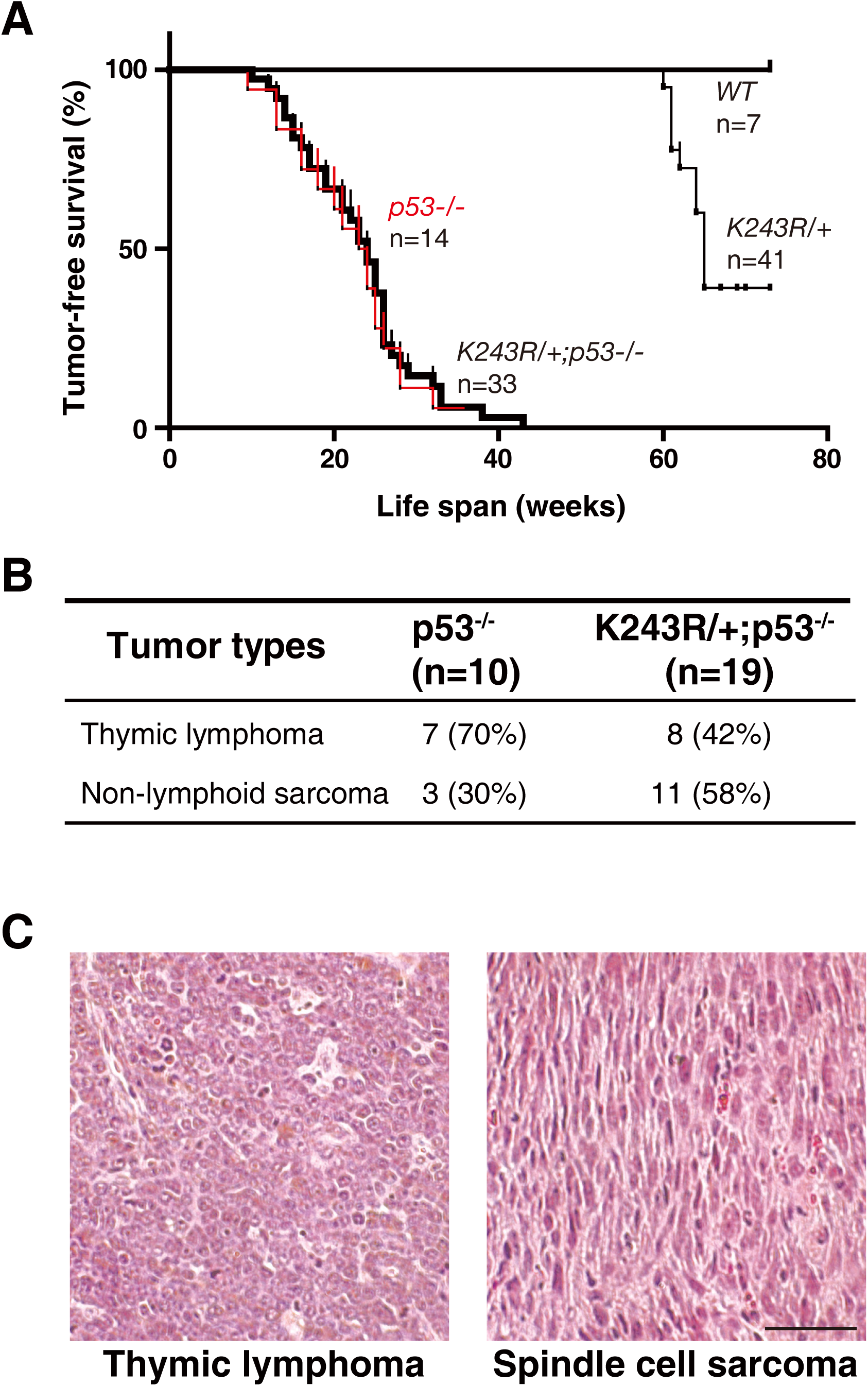
Intercross with *p53*-deficient (*p53-/-*) mice leads to the acceleration of tumorigenesis and change in tumor spectrum in acetylation-deficient BubR1 (*K243R/+*) mice. **(A)** Tumor free survival (log-rank test) of wild-type (*WT), K243R/+, p53-/-*, and *K243R/+*; *p53-/-* mice, respectively. Tumor incidence was scored by histologic examination of tissues from sacrificed mice. The median tumor-free survival time was scored and marked. n, number of mice examined; *K243R/*+ & *K243R/+; p53-/-, p* < 0.0001; p53-/-& *K243R/+;p53-/-, p* = 0.8282 (log-rank test). **(B)** Spectrum of tumors from *p53-/-* and *K243R/+*; *p53-/-* mice, respectively. **(C)** H & E-staining of lymphoma and spindle cell sarcoma found from *K243R/+*; *p53-/-* mice. Scale bar, 50 μm.

Interestingly, the tumor spectrum of *K243R/+; p53-/-* mice differed from that of *K243R/+* mice. Tumors of *K243R/+* mice do not show tissue specificity: most tumors were of mesenchymal origin, such as lymphomas, sarcomas, hepatocellular carcinomas (11). However, we noticed an interesting change of spectrum when *K243R/+* was crossed to *p53-/-* mice. It is well established that tumors of *p53*-/- mice are mostly thymic lymphomas and some sarcomas (17, 18). Results were recapitulated in our breeding scheme that tumors of *p53*-/- mice were 70% thymic lymphoma (n=7) and 30% sarcomas (n=3). Interestingly, tumors from *K243R/+; p53-/-* mice exhibited similar pattern to *p53-/-* that tumors were mostly sarcomas and lymphomas but with a shift in tumor spectrum: 58 % non-lymphoid sarcomas (n=11) and 42 % thymic lymphomas (n=8) (Fig. 1B). Lymphoma or non-lymphoid sarcoma was disseminated to multiple sites, including thymus, leg, back, or bone (data not shown). Examples of thymic lymphoma and spindle cell sarcoma from *K243R/+; p53-/-* mice are shown (Fig. 1C). Collectively, intercross to *p53-/-* led to shortened tumor latency with the change in tumor spectrum from that of *K243R/+* mice.

### Loss of BubR1 acetylation induces replication stress as well as PMSCS and aneuploidy

Near diploid aneuploidy and premature sister chromatid separation (PMSCS) are hallmarks of *K243R/+* mice (11). Numerous studies report polyploidy and aneuploidy in *p53*-deficient mice, which contribute to tumorigenesis (19). Indeed, our analysis corroborated that *p53-/-* MEFs exhibit a high incidence of aneuploidy and polyploidy, whereas *K243R/+* MEFs display aneuploidy but not polyploidy. In *K243R/+; p53-/-* MEFs, aneuploidy was apparent (Fig. 2A).

**FIG 2.**
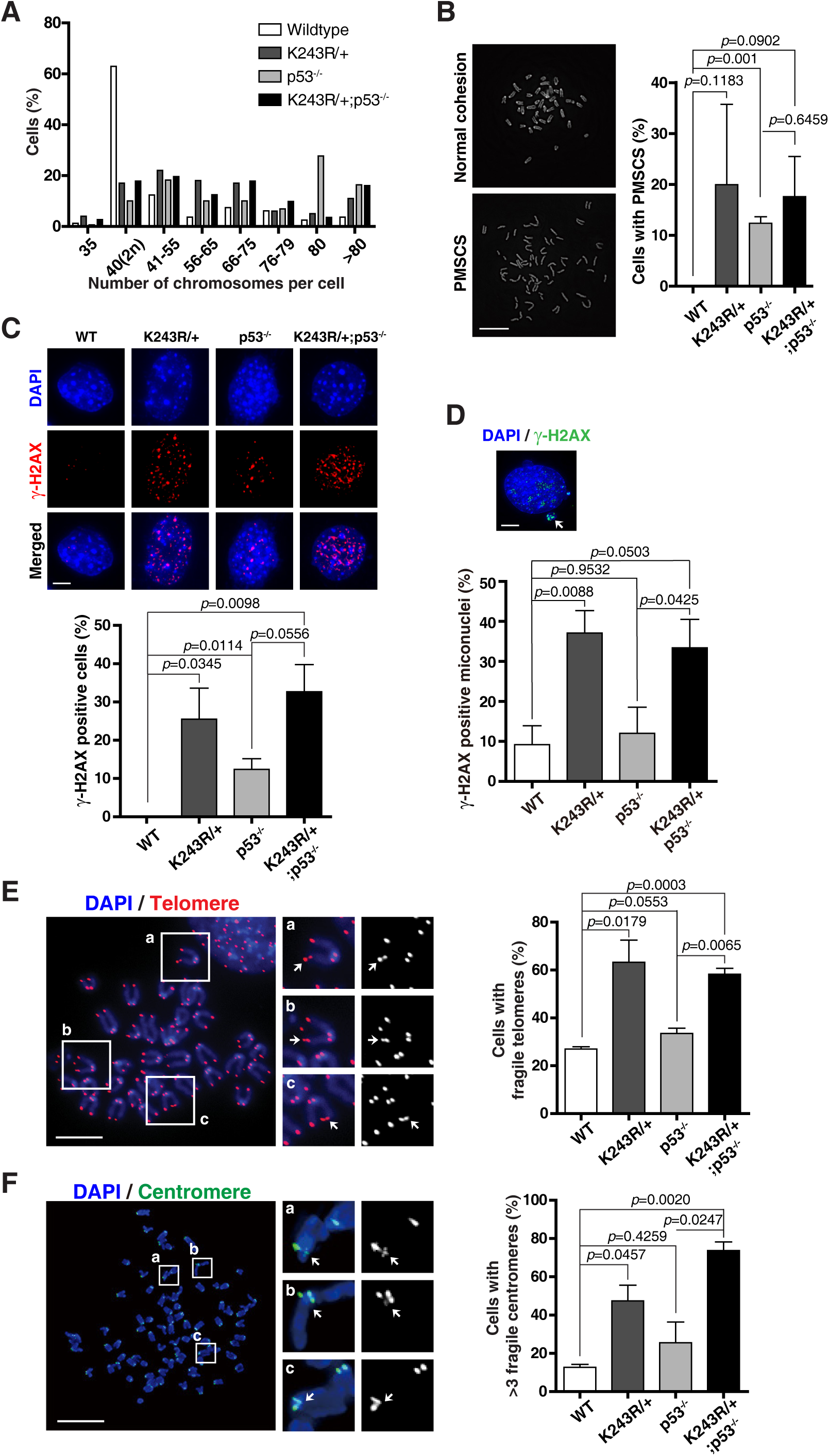
Aneuploidy, global DNA damage, and fragile telomeres and centromeres in K243R/+;*p53-/-* mice. **(A)** The fraction of MEFs in the indicated number of chromosomes. Fifty cells each from the indicated group were scored. Total of 2000∼ 2500 chromosomes was counted in each group. **(B)** Premature sister chromatid separation (PMSCS) in MEFs. DAPI stained chromosomes were assessed for PMSCS (right). Representative images of usual cohesion and PMSCS are shown (left). Twenty chromosome spreads from each group were scored in three independent experiments (mean ± SEM; n = 60). Scale bar, 10 μm. *P* values were obtained from the student’s t-test. **(C)** MEFs were subjected to immunofluorescence with anti-γ-H2AX antibodies, followed by counterstaining with DAPI. Scale bar, 5μm (upper panel). Number of cells with γ-H2AX foci were counted and depicted as bar graphs (lower panel). Twenty cells from each group were scored in three independent experiments (mean ± SEM; n = 60). *P* values were obtained from the student’s t-test. **(D)** The frequency of micronuclei co-localized with γ-H2AX signal was counted (lower panel). Twenty cells from each genotype were analyzed in three independent experiments. (mean ± SEM; n = 60) *P* values were obtained from the student’s t-test. Representative image of γ-H2AX positive micronuclei is shown. Scale bar, 5 μm (upper panel). **(E)** Telomere-FISH on metaphase chromosome spreads reveal replication stress. Enlarged images of the insets are shown. Arrows mark discontinuous gapped FISH signals (left). Frequency of cells with fragile telomeres in each group (right). Bars are the result of three independent experiments (mean ± SEM; n = 60). *P* values were obtained from the student’s t-test. Scale bar, 5 μm. **(F)** Centromere integrity was examined by centromere-FISH. Enlarged images of the insets are shown. Arrows mark fragile centromeres (left). Frequency of cells with fragile centromeres (right). Twenty chromosome spreads from each group were scored in three independent experiments (mean ± SEM; n = 60). *P* values were obtained from the student’s t-test. Scale bar, 10 μm.

PMSCS, which results from premature activation of APC/C without the satisfaction of SAC, was over 25% in *K243R/+* MEFs and ∼ 10% in *p53*-deficient MEFs. The incidence of PMSCS in *K243R/+; p53-/-* was not additive but were similar to that of *K243R/+* MEFs (Fig. 2B). Taken together, PMSCS in *K243R/+; p53-/-* resembles *K243R/+*, and less to *p53* deficiency.

When *p53* cDNA was sequenced from the primary tumors of *K243R/+* mice, it showed many missense mutations that were reported oncogenic (11). The result suggested to us that genetic instability is triggered by the loss of BubR1 acetylation. Therefore, we asked if DNA damage increased by the loss of BubR1 acetylation.

Immunofluorescence with anti-γ-H2AX antibodies revealed that spontaneous DNA damage was apparent in *K243R/+* MEFs, even more than *p53*-null. In *K243R/+; p53-/-* MEFs, DNA damage markedly increased, indicative of an additive effect of *p53* deficiency and the loss of BubR1 acetylation (Fig. 2C). However, γ-H2AX-positive micronuclei, which result from chromosome mis-segregation (12), were markedly increased in *K243R/+* and *K243R/+; p53-/-* MEFs, but significantly lower in *p53-/-* MEFs (Fig. 2D). These results suggest that loss of BubR1 acetylation leads to DNA damage, and the underlying mechanism differs from that of *p53* deficiency.

Immunofluorescence with anti-γ-H2AX immunostaining, coupled with telomere- or centromere-FISH were performed to assess if telomeres and centromeres were affected. In interphase, both telomeres and centromeres were damaged in *K243R/+* and *p53-/-*, respectively, which worsened in *K243R/+; p53-/-* MEFs (Supplemental Fig. S1A and B).

Due to the tandem repeat sequences, centromeres and telomeres are hard-to-replicate regions that can impede the progression of replication forks, eventually leading to the fragility in the event of replication stress (20). We analyzed the centromere and telomere fragility by employing telomere- and centromere-Fluorescence in situ hybridization (FISH) on metaphase chromosome spreads to accurately assess the fragility. FISH signals that were not forming punctate foci but gapped or tailed were scored fragile (Fig. 2E and F, arrows in inset). The result showed that *p53*-null MEFs exhibited a slight increase in centromere and telomere fragility, but the fragility was markedly increased in *K243R/+* and *K243R/+; p53-/-* MEFs both in centromeres and telomeres. Notably, telomere fragility was far more affected, when compared to centromeres (Fig. 2E and F, graph). These data altogether suggest that loss of BubR1 acetylation led to replication stress and that a significant portion of DNA damage in *K243R/+* come from the dramatic increase of the replication stress.

### Tumorigenesis by the loss of BubR1 acetylation involves various chromosome structural aberrations

Errors in mitosis and its resultant micronuclei were suggested to be the sources of chromothripsis (12, 13), one-off catastrophic chromosome breakage, and rearrangement in cis (14). Chromothripsis is the cause, rather than the result, of aggressive cancers of mesenchymal origin. As tumorigenesis by the loss of BubR1 acetylation in *K243R/+* result from mitotic infidelity, we asked if chromothripsis could result from BubR1 acetylation deficiency.

Next-generation sequencing of the whole genome was performed for the primary liver cancer obtained from *K243R/+*, cultured sarcoma cells of *p53-/-*, and sarcoma cells of *K243R/+; p53-/-*, respectively. Deep sequencing was performed at an average depth of ∼ 41X, alongside with the paired constitutional DNA from healthy liver tissue. Then the sequencing data were analyzed for the copy number changes, using BIC-seq2 (21).

Using Meerkat followed by specific quality filtering (22), we found 11 predicted inter- and intra-chromosomal breakpoints in the cultured sarcoma cells of *p53-/-* mice. In comparison, there was a marked increase of intra- and inter-chromosomal breakpoints and rearrangements by the loss of BubR1 acetylation: 113 breakpoints in *K243R/+* primary liver cancer and 127 breakpoints in *K243R/+;p53-/-* sarcoma cells, respectively (Fig. 3A). The fidelity of predicted breakpoints was confirmed with PCR validation (data not shown).

**FIG 3.**
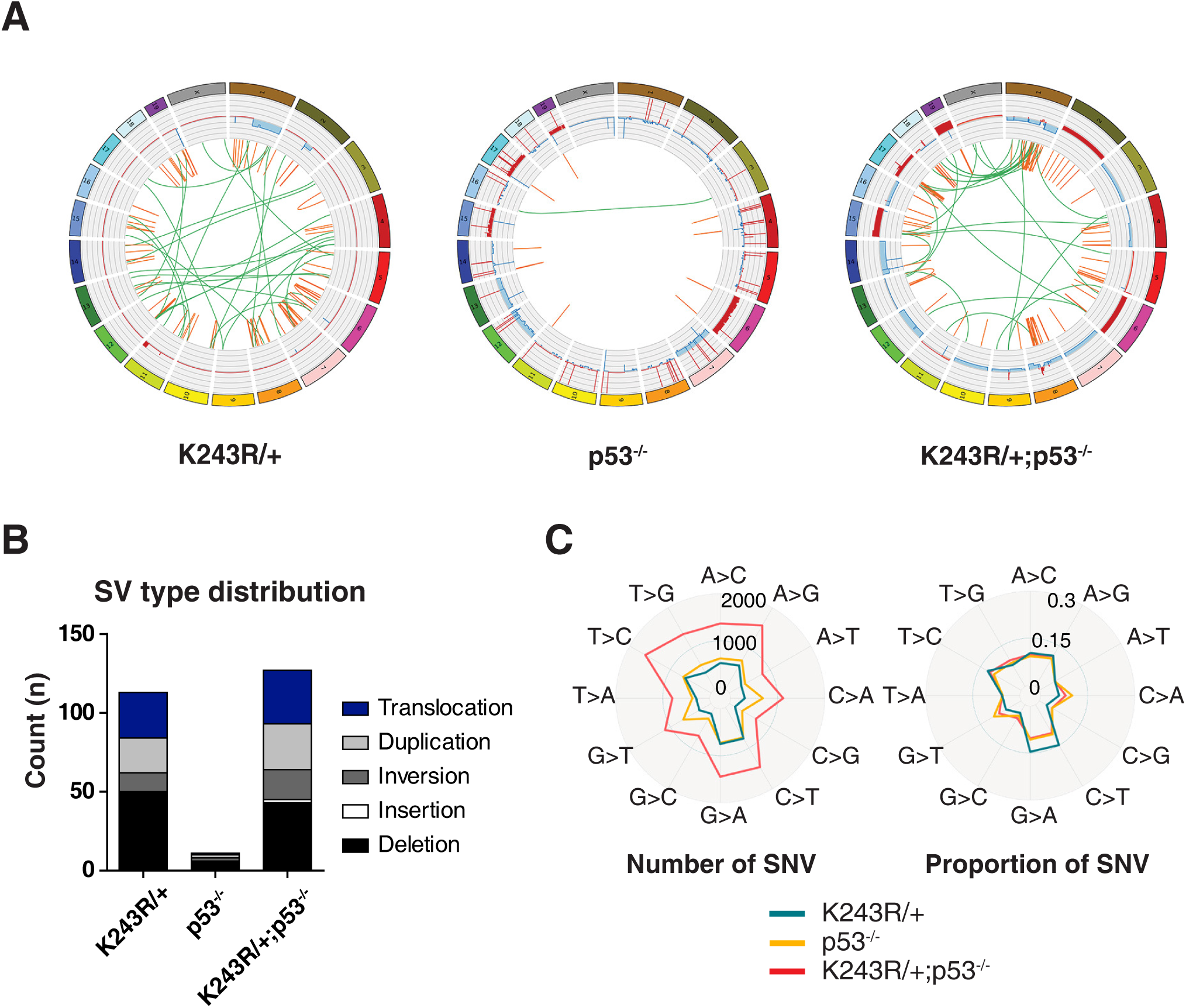
Whole-genome sequencing and analysis reveal frequent intra- and inter-chromosomal translocation by the loss of BubR1 acetylation. **(A)** Circos map of whole-genome sequencing. Data are from primary liver cancer of *K243R/+*; cultured sarcoma from *p53-/-*; cultured sarcoma from *K243R/+;p53-/-* mice. Chromosomes are individually colored. Copy numbers of the genomic region are plotted as gains in red and losses in blue, using log2 ratio. Chromosomal rearrangements are depicted on each side of the breakpoints; intra-chromosomal rearrangements (orange) and inter-chromosomal rearrangements (green) are located in the center of the circle. **(B)** Chromosome structure variation (SV) in tumors of *K243R/+, p53-/-,* and *K243R/+; p53-/-*. Frequency of translocation, duplication, inversion, insertion, and deletion are depicted as bar graphs. **(C)** The spectrum of single nucleotide variation (SNV) from each tumor cells. Total numbers (left) and the proportions of somatic SNVs (right) are shown respectively.

One of the characteristics of chromothripsis is the oscillation between two copy number states (23). In our analysis, the oscillation between two copy number states was hard to define; instead, complex copy number variations were apparent (Fig. 3A). It is possible that due to the complex rearrangements of various types, chromothripsis could not be easily verified. Chromosome duplication, inversion, insertion, and deletion were also markedly induced by the loss of BubR1 acetylation (Fig. 3B). Taken together, these data suggested to us that loss of BubR1 acetylation resulted in chromosome shattering that led to complex structural variations, such as intra- and inter-chromosomal translocation, rearrangements, and potentially chromothripsis.

The analysis of genome-wide single-nucleotide variant (SNV) spectrum revealed predominant mutation type with T>C, A>G, G>A, and C>T changes being most common, followed by T>G and A>C (Fig. 3C). This spectrum was broadly similar among *K243R/+, p53-/-*, and *K243R/+;p53-/-* tumor cells. The total numbers of SNVs were 5,140 in *K243R/+* tumor cells, 6,801 in *p53-/-* cells, and 14,022 in *K243R/+;p53-/-* cells, respectively. (Fig. 3C, left). However, G>T and C>A mutations were relatively frequent in *p53-/-* cells, and T>C mutations were most frequent in *K243R/+* tumor cells (Fig. 3C, right). The results suggest that genetic instability, reflected by random point mutations accompany the loss of BubR1 acetylation in tumorigenesis.

### Robertsonian whole-arm translocation is significant in tumors of BubR1 acetylation-deficient mice

As centromere and telomere were damaged by the loss of BubR1 acetylation in primary MEFs, we asked whether tumors from *K243R/+; p53-/-* exhibit any further chromosome aberrations related to centromeres and telomeres. When centromere and telomere-FISH were performed in tumor cells, we made an interesting observation: most apparent chromosome aberration was the Robertsonian-like whole-arm translocation that led to X-shaped chromosomes instead of U-shaped telocentric mouse chromosomes in *K243R/+; p53-/-* cells (Fig. 4A). Whole-arm translocation was a significant feature in tumors of *K243R/+; p53-/-*, but was rarely found in *p53-/-* tumors (Fig. 4A, right).

**FIG 4.**
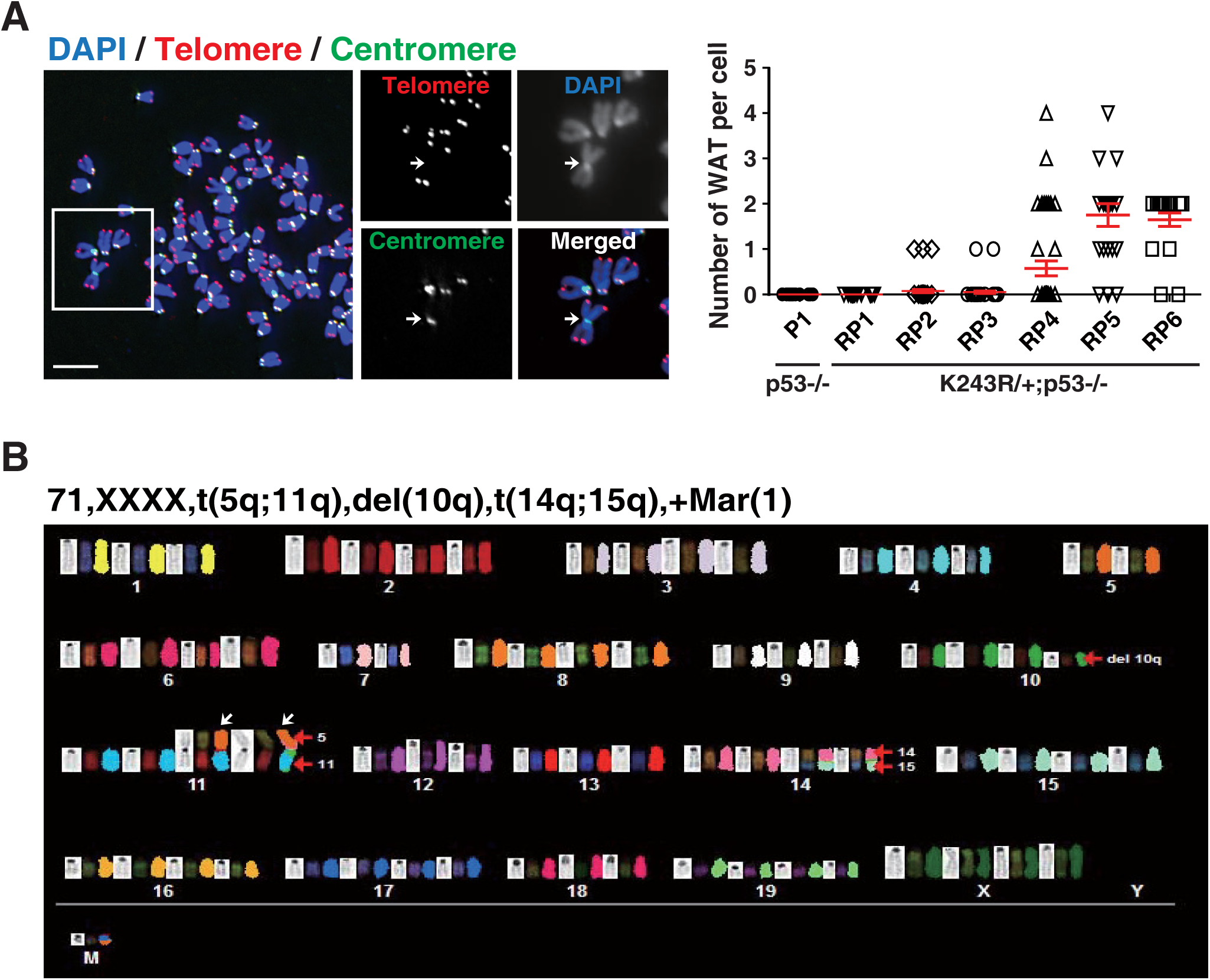
Chromosomes from Tumors of *K243R/+; p53-/-* display Robertsonian whole-arm translocation. **(A)** Metaphase chromosomes of established sarcoma and lymphoma cell lines from *K243R/+;p53-/-* mice were subjected to centromere and telomere-FISH. Centromeres were detected by hybridization with FAM-labeled probe (green), and telomeres were hybridized with Cy3-labeled probe (red). Chromosomal DNA was counter-stained with DAPI (blue). Arrows mark whole-arm translocation (WAT) without telomeres at the fusion points. Scale bar, 5 μm (left). Number of whole-arm translocations per cell is depicted. P1, *p53*-deficient sarcoma; RP1-6, tumors from *K243R/+; p53-/-* (RP1-4, sarcomas; RP5 & RP6, lymphomas). All tumor cells were analyzed at early passage (below passage 10 from primary tumors). Forty cells each were scored (mean ± SEM, n=40) (right). **(B)** Representative spectral karyotyping of a sarcoma (RP4) from *K243R/+;p53-/-* mice. White arrows mark whole-arm translocations. Karyotype of the examined tumor cell is indicated.

The fusion of two chromosomes at the short arm, but not the long arm, where centromeres and telomeres are proximal to each other generated X-shaped chromosome. At the fused point, telomere signal was lost, whereas centromere was present (Fig. 4A, arrows in enlarged images). This suggests that telomeres were eroded inducing end-end fusions, while centromere specification was unperturbed in the fused chromosomes. Consistently, telomere and centromere fragility were apparent, as were in MEFs (Fig. 4A, arrows in the inset).

Robertsonian whole-arm translocation was a significant feature of the tumors from *K243R/+; p53-/-* but was rarely detected from *p53-/-* tumors (Fig 4A, graph). These results suggest that the replication stress at the telomeres, caused by the loss of BubR1 acetylation may be the underlying reason for telomere erosion and subsequent whole-arm translocation. Spectral karyotyping of the tumor cells corroborated the whole-arm translocation in *K243R/+; p53-/-* sarcoma cells, as well as inter-chromosome rearrangements (Fig. 4B, arrows). The fusion between chromosomes was random and not restricted to specific chromosomes, at least at this stage of culture (passage 30), although the example in the figure shows the translocation between chromosomes 5 and 11 (Fig. 4B).

### Loss of BubR1 acetylation is found in human cancers

BubR1 acetylation deficiency in mice leads to tumorigenesis through chromosome mis-segregation, due to weakened SAC and failure in the stabilization of chromosome-spindle attachment (11). We further showed here that BubR1 acetylation deficiency results in various types of chromosome structural aberration due to replication stress.

We next asked whether BubR1 acetylation deficiency could be found in human cancers. In the cancer genome database (cBioPortal, TCGA), we did not find the exact mutation of K250 in *BubR1*. However, we found nine mutations around K250 of *BubR1* from the TCGA database using cBioPortal (Supplemental Table S1) (24-28).

We tested some of the mutants for their capability of acetylation of BubR1, using the monoclonal antibody specific to BubR1 acetylated at K250. Four mutants (Fig. 5A, bottom) that were tested for the capability of BubR1 acetylation are as follows: *BubR1* mutation at *R244C* and *G246R*, reported from melanomas; *A251P* mutation from renal cell carcinoma; *N255Y* mutation from Rhabdomyosarcoma (29, 30) (Supplemental Table S1). Mutant *BubR1* constructs were generated by site-directed mutagenesis, with Myc tagged at the N-terminus then were transfected into 293T cells. Two days post-transfection, cells were treated with nocodazole to arrest in mitosis, then subjected to immunoprecipitation (IP) with anti-BubR1 antibodies, followed by Western blot (WB) analysis with monoclonal anti-acetyl-K250 antibody (7, 8). For control, wild-type *BubR1* (WT) and acetylation-deficient mutant *BubR1* (K250R)-expressing constructs were employed.

**FIG 5.**
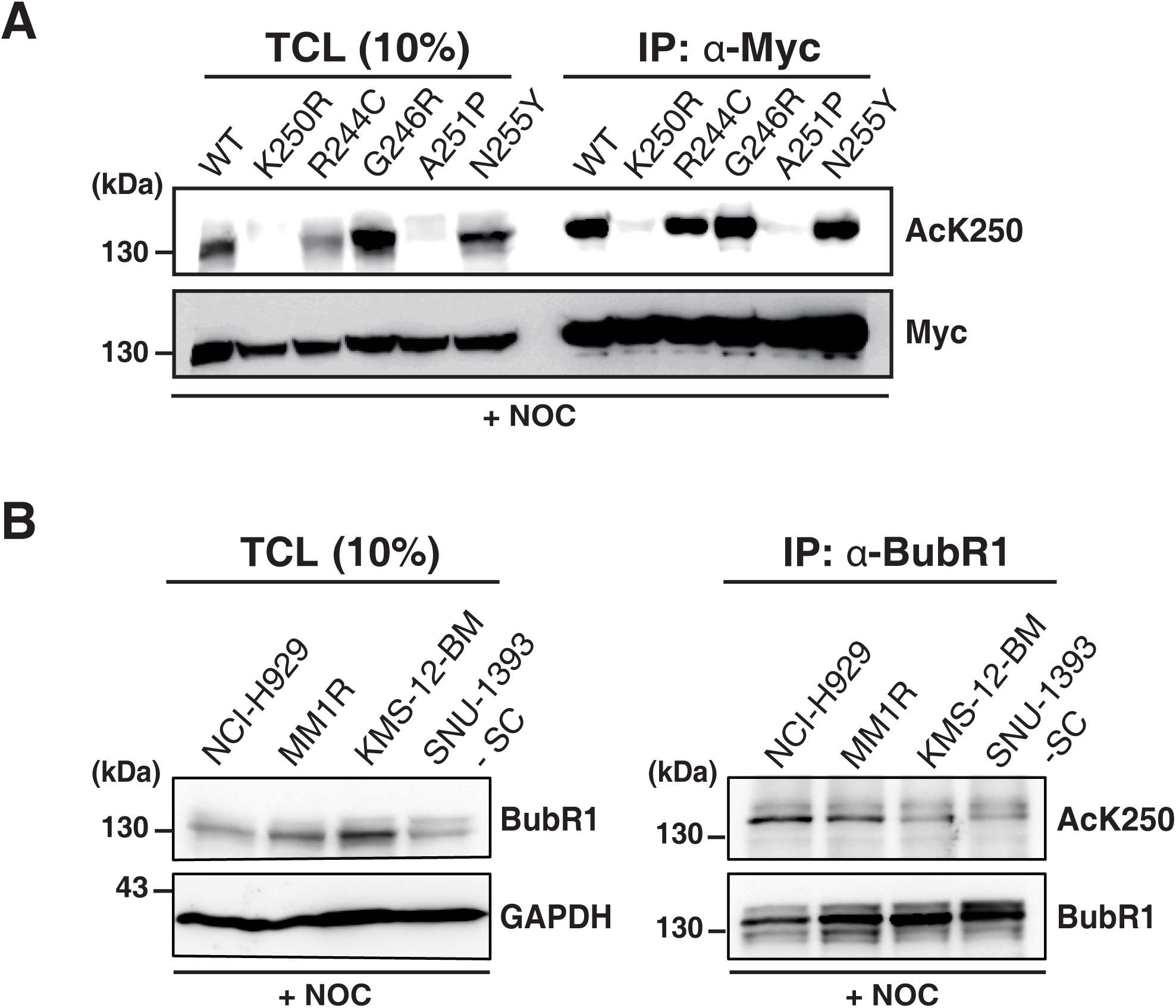
BubR1 acetylation deficiency is found in human malignant tumors. **(A)** Various BubR1 mutations adjacent to K250 acetylation site were selected from the database of The Cancer Genome Atlas (TCGA). Site-directed mutagenesis was performed on *BubR1*-expressing construct, tagged with Myc. Then the constructs were transfected to 293FT cells, followed by nocodazole treatment to enrich cells in mitosis. The selected mutations were examined for the capability of BubR1 acetylation by immunoprecipitation (IP) with anti-BubR1 antibody, followed by Western Blot (WB) analysis using monoclonal antibody against acetylated BubR1 (anti-AcK250). Same blot was reprobed with anti-Myc (9E10) for loading control. Ten percent of total cell lysates (TCL) were subjected to WB for control. K250R mutant was employed as a control. **(B)** Human multiple myeloma cell lines NCI-H929, MM1R, KMS-12-BM and SNU-1393-SC were assessed for the capability of BubR1 acetylation in mitosis by IP and WB, using anti-AcK250. Cells were arrested in mitosis by nocodazole treatment. Ten percent of total cell lysates (TCL) were subjected to WB analysis.

The result showed that *A251P* mutant, which was found from renal cell carcinoma was completely defective in BubR1 acetylation, as was the control *K250R* (Fig. 5A). In total cell lysates, *R244C* mutant found from melanoma displayed a markedly lower level of BubR1 acetylation when compared to the normalized protein expression level (Fig. 5A, anti-Myc, bottom). These results indicate that BubR1 acetylation deficiency is associated with human cancers.

Lysine 250 of BubR1 resides at the unstructured region near to D-box, ABBA motif, and KEN motif, which are the motifs critical in recognizing APC/C and/or Cdc20 (3, 4, 9). As this region is crucial in modulating APC/C, and the structure of APC/C-MCC is complex, one cannot merely assume the capability of acetylation before testing. Here, one out of four mutants tested showed complete loss of BubR1 acetylation, and one partial loss, in mitosis. We expect that the list of BubR1 mutations that turn out to be defective in acetylation at K250 in mitosis will increase. Furthermore, it should be noted that the tumor suppressor BRCA2 is needed for BubR1 acetylation, recruiting PCAF acetyltransferase to BubR1 (31). Therefore, *BRCA2*-deficient cells exhibit defective BubR1 acetylation/deacetylation and checkpoint signaling (8) even without the mutation of *BubR1* (31). Collectively, BubR1 acetylation deficiency is associated with human tumorigenesis either by a direct mutation in *BubR1* or due to defective *BRCA2*.

Next, we asked if whole-arm translocation is also a feature associated with BubR1 acetylation deficiency in human cancer. Robertsonian-like whole-arm translocation in human somatic cells is rare, probably due to the fact that human chromosomes are not telocentric. In the case where whole-arm translocations are, they are usually at the acrocentric chromosomes (32). However, cytogenetic analysis of hematopoietic malignancies in Seoul National University Hospital (SNUH) showed that 83 cases out of 1,837 patients display whole-arm translocation: multiple myeloma being the most frequent type (11.6%) (Table 1).

**TABLE 1.** Frequency of whole-arm translocation in hematopoietic malignancies from patients of Seoul National University Hospital (SNUH)

| Disease | Number of patients | Cases diagnosed |
| --- | --- | --- |
| Acute myeloid leukemia | 730 | 15 |
| Multiple myeloma | 334 | 39 |
| Acute lymphoblastic leukemia | 219 | 2 |
| Myelodysplastic neoplasms | 158 | 13 |
| Lymphoma | 128 | 5 |
| Diffuse large B-cell lymphoma | 80 | 1 |
| Chronic myeloid leukemia | 79 | 1 |
| Aplastic anemia | 49 | 1 |
| Chronic lymphocytic leukemia | 23 | 1 |
| Essential thrombocythemia | 23 | 2 |
| Idiopathic myelofibrosis | 11 | 1 |
| Monoclonal gammopathy of undetermined significance | 2 | 1 |
| Myelomatosis | 1 | 1 |
| Total | 1837 | 83 |

Having these information into account, we searched for multiple myeloma cell lines that displayed whole-arm translocation (WAT) to ask if BubR1 acetylation deficiency is associated with WAT in human malignancies. Four cells lines were selected for IP and WB: two multiple myeloma cell lines KMS-12-BM (33) and SNU-1393-SC (34) where we could find complete cytogenetics including WAT; NCI-H929 that was negative for WAT (35); MM.1R cell line, which lacks complete cytogenetic information (36) (Supplemental Table S2).

Cells were treated with nocodazole, enriched for mitotic cells, then subjected to IP with anti-BubR1 antibodies and WB with monoclonal anti-acetyl-K250 antibody (8). Interestingly, the result showed that Robertsonian whole-arm translocation may be linked with BubR1 acetylation deficiency: two cell lines that were reported for whole-arm translocation, KMS-12-BM and SNU-1393-SC, displayed partial loss of BubR1 acetylation, whereas NCI-H929 and MM.1R exhibited BubR1 acetylation in this setting (Fig. 5B, right). Level of BubR1 in total cell lysate is shown for comparison (Fig. 5B, left).

## Discussion

Aneuploidy, PMSCS, missense mutation, micronuclei, and chromosome translocation were also found from tumors in *K243R/+* mice (11). It suggested to us that the loss of BubR1 acetylation might cause various chromosome alterations. To address this issue, rescue of all chromosome defects was needed. By interbreeding *K243R/+* mice to *p53*-null mice and generate double mutant mice (*K243R/+; p53-/-*), we achieved this aim. The generation of the double mutant mice was physiologically supported by the fact that 8 out of 11 primary tumors of *K243R/+* mice displayed *p53* missense mutations (11). In this study, we found that many features of *K243R/+; p53-/-* followed *K243R/+*, rather than *p53-/-* in chromosome instability.

We learned several unexpected things from this breeding scheme. First, breeding of *K243R/+* mice with *p53-/-* mice changes the tumor spectrum: decrease of thymic lymphoma and increase in sarcoma, compared to *p53-/-* (Fig. 1B). Interesting though, we do not understand the reason for the change in spectrum.

Secondly, we have identified that the loss of BubR1 acetylation results in fragility of heterochromatin. Fragility in telomeres and centromeres result from failure of replication fork progression (20), also called replication stress. BubR1 acetylation deficiency, at first sight, seemed unrelated to the replication stress. It was reported that aneuploidy can cause stress in replication, due to the imbalance of replication machinery and the amount of DNA to be replicated (37, 38). However, *p53*-deficient MEFs exhibited markedly less replication stress at centromeres and telomeres, albeit aneuploidy and polyploidy, compared to *K243R/+*. Therefore, the replication stress observed in *K243R/+* and *K243R/+; p53-/-*MEFs may be a more direct result of the loss of BubR1 acetylation than from aneuploidy. Interestingly, a recent report showed that Bub1-Bub3, components of the mitotic checkpoint that interact with BubR1, promotes telomere replication in S phase (39). Therefore, there is a possibility that the defect in BubR1 acetylation may have affected the integrity of Bub3-Bub1 in S phase telomere replication fidelity. Alternatively, it could be that mitotic infidelity generally affects replication integrity like the case in yeast (40).

Mechanism of centromere establishment and specification are very different from those of telomeres (41, 42). Centromeres do not require specific nucleotide sequences but is specified by the loading of centromere specification proteins. In comparison, telomeres are consisted of (TTAGGG)n repeats and are capped through the formation of telomere-loops (T-loops). If not protected, telomeres are exposed to DNA damage response pathway, inducing repair and recombination pathway (43). Therefore, the result demonstrated here suggest that deficiency of BubR1 acetylation, which takes place at the centromeric kinetochore, affects the integrity of telomeres, suggesting that there is a crosstalk between centromeres and telomeres for chromosome integrity.

We have observed end-end fusion at the short arms, generating whole-arm translocation in tumors of *K243R/+; p53-/-*. Like this case, it is reported that dicentric chromosomes generated by telomere end fusions can result in the chromosome shattering. Furthermore, telomere crisis are associated with chromothripsis and kataegis, leading to tumorigenesis (44). It would be interesting to see if kataegis can be observed in tumors by the loss of BubR1 acetylation. Of note, there is a possibility that *BRCA2*-deficiency may be associated with kataegis (45).

We were not able to detect chromothripsis, probably due to the complex structural aberrations in tumors of *K243R/+*. Instead but interestingly, we found Robertsonian-like whole-arm translocation in tumors derived from the loss of BubR1 acetylation (*K243R/+; p53-/-*). We interpreted that this is due to the telomere end-end fusions derived from replication stress. Mouse chromosomes are telocentric, with centromere and telomere in close proximity at the short arm, the Robertsonian whole-arm rearrangements are probably more frequent in mouse chromosomes than in human. In human malignancies (46), the chromosome that exhibited whole-arm translocation are not random but are found at acrocentric chromosomes (32). Nonetheless, we observed a possibility that defects in BubR1 acetylation in mitosis might be linked with complex chromosome rearrangements including whole-arm translocation in a subset of human multiple myeloma cell lines (Fig. 5B).

Moreover, genome-wide sequencing and spectral karyotyping revealed a significant increase of inter- and intra-chromosome rearrangements and various types of genetic instability. These massive genetic alterations are the cause of high incidence tumorigenesis by the loss of BubR1 acetylation. Aneuploidy, in addition, would have aggravated the situation rather than being the primary cause of tumorigenesis.

Whether BubR1 K250 mutation or acetylation deficiency can be found from human cancers was an earnest question, although *BRCA2*-deficient cancers exhibit defective BubR1 acetylation (31). Using the public cancer genome atlas, TCGA, we were able to pull out 9 mutations proximal to K250. (Supplemental Table S1). Out of four tested, *A251P* mutation showed a complete defect in BubR1 acetylation at K250, as is in *K250R*, and one reduced level of BubR1 acetylation (R244C). These results indicate that defects in BubR1 acetylation is likely to be associated with human cancers.

Forced expression of the acetylation-mimetic form of BubR1 (K250Q) led to apoptosis in *BRCA2*-deficient cells (10). Furthermore, *BRCA2*-deficient cells exhibited resistance to pan-HDAC inhibitors because they lack BubR1 acetylation, but was sensitive to a specific type of HDAC inhibitor (8). It would be interesting to see if cancers with defective BubR1 acetylation will be sensitive to K250Q-BubR1 expression or a specific type of deacetylase inhibitors.

Previously, we have seen that BubR1 level was associated with poor prognostics of recurrence-free survival of ovarian cancers after surgery (47). The results shown here suggest that assessing BubR1 acetylation by utilizing the acetylated BubR1-specific antibody may reveal clinical values. Targeting BubR1 acetylation pathway may be an effective treatment for cancers with *BRCA2* mutation and also cancers of chromosome instability, particularly the ones with complex chromosome rearrangements.

## Materials and Methods

### Statistical analysis

The probability of tumor-free survival was estimated using the Kaplan-Meier method and analyzed by the log-rank test. For the other statistical analyses, t-test was used unless otherwise stated. The statistical data were analyzed using GraphPad Prism5.

### Plasmid construction and transfection

Various BubR1 mutants were generated by site-directed mutagenesis using pcDNA3.1-myc-BubR1 as the template. Transfections of BubR1 mutant constructs into HEK293 cells were performed using Lipofectamine 2000 (Invitrogen).

### Animal care, approval, and IRB approval of human cancer data

*K243R/+* mice were previously reported (11). *K243R/+* mice were crossed with *p53 +/-* mice (B6.Cg-Trp53^tm1Ty^/J) several times to obtain *K243R/+; p53-/-* mice. Mice used in this study are B6;129 mixed background. Mice were housed in a specific pathogen-free facility. The Institutional Animal Care and Use Committee of Seoul National University (SNU-151008-3-5) approved the experimental animal protocols. We followed the guidelines and policies for the Care and Use of Laboratory Animals at Seoul National University. Karyotype analysis of hematologic malignancies from Seoul National University Hospital (SNUH) was approved from the IRB of SNUH (1103-004-353).

### Histopathology

Tissues were harvested from the following organs: thymus, lung, liver, spleen, skin, spine, and lymph nodes. Tissues were fixed in 4% formaldehyde, embedded in paraffin, and sectioned. The sections were deparaffinized in xylene, rehydrated in ethanol, and subjected to hematoxylin and eosin (H&E) staining. Two different pathologists made the diagnosis of the diseases by analyzing the H&E stained specimens (Logone Bio Convergence Research Foundation, Korea).

### Detection of somatic copy number alterations

Somatic copy number alterations (SCNAs) were identified using the BIC-seq2 (version 0.6.3) (21). BIC-seq2 algorithm is composed of two main components. Firstly, we normalized potential biases in the aligned sequencing reads data, based on the mouse reference genome (GRCm38), at the nucleotide level with 1000 bp binning parameter. Secondly, SCNAs were detected based on Bayesian information criterion-based segmentation. We used three as the value of lambda parameter, which is the main parameter defines the smoothness of the SCNAs profile.

### Detection of structural variations

Identification of structural variations (SVs) was conducted using the Meerkat (version 0.189) (22). Sequencing reads were aligned against the mouse reference genome (GRCm38) to identify the soft-clipped and unmapped reads. These reads were remapped to identify discordant read pairs. After adjusting non-uniquely mapped reads, we paired clusters to call multiple events. Candidates of breakpoint regions were predicted by supporting reads, and precise breakpoints were identified by local alignments. Somatic events were obtained by further processes filtering out germline and low confidence calls.

### Detection of somatic single nucleotide variants and short indels

We used MuTect2 included in the Genome Analysis Toolkit (version 3.5) with default parameters to detect somatic single nucleotide variations (SNVs) and short indels (48). MuTect2 is a method that applies a Bayesian classifier to identify somatic SNVs and short indels with low allele fractions followed by careful filtering processes allowing high specificity of each mutation.

### Cell culture

Multiple myeloma cell SNU-1393-SC was previously established (34). NCI-H929, MM1R, and KMS-12-BM were obtained from either ATCC or DSMZ. HEK293T cells and mouse tumor cells were cultured in DMEM, supplemented with 10% Fetal Bovine Serum (Lonza, Switzerland).

### Metaphase spreads and Fluorescence *in situ* hybridization (FISH)

Metaphase chromosome spreads and FISH were performed according to the previous protocol (49). Images were acquired with a microscope (DeltaVision; Applied Precision) equipped with 60X objective lens (Olympus). Then the images were obtained with 0.2-μm-distanced optical sections in the z-axis. Each section was deconvoluted and projected, using the softWoRx software (Applied Precision).

### Immunofluorescence assay

Immunofluorescence was performed following the previous publication with slight modifications (7, 8, 11, 31). Antibodies used were as follows: anti-phospho-histone H2A.X antibody (Ser 139; Millipore); anti-human ACAs (CREST; Antibodies Incorporated).

### Spectral karyotyping assay

For the spectral karyotyping assay, tumor cells from *K243R/+; p53-/-* mouse were treated with 5 μg·ml^-1^ colcemid for 2 h, followed by fixation. The spectral karyotyping assay was performed at the Molecular Cytogenetics Core at the University of Texas M.D. Anderson Cancer Center.

### Immunoprecipitation (IP) and Western blot (WB) analysis

For enrichment of mitotic cells, HEK293 or Multiple myeloma cells were treated for 18h with 200 ng·ml^-1^ nocodazole (Sigma-Aldrich). IP and WB followed the previous protocol (7, 8) with the antibodies listed: anti-c-myc (sc-40; Santa Cruz Biotechnology); anti-human BubR1 (612503; BD biosciences); anti-GAPDH (AbC-1001; AbClon) antibodies. Home-made monoclonal antibody against AcK250 BubR1 was previously described (8).

## Acknowledgment

This work was supported by Tissue Regeneration Research (2016M3A9B4918405) and Post-genome Research (2017M3C9A5030991, 2017M3C9A5031002 and 2017M3C9A5031004) from the Korean Research Foundation. We are grateful to Si-Young Choi for assisting the experiments. We thank all members of the Lee lab for helpful discussions throughout the study. The results here are in part based upon data generated by the TCGA Research Network: https://www.cancer.gov/tcga.

## Authors’ contributions

HL initiated the project and wrote the manuscript; JP and SY conceived, designed and performed experiments; SP, JL performed experiments in chromosome analysis; JH and SL performed genomic analysis; YK collected patient data and is responsible for IRB approval.

